# Adversarial manipulation of human decision-making

**DOI:** 10.1101/2020.03.15.992875

**Authors:** Amir Dezfouli, Richard Nock, Peter Dayan

**Affiliations:** Data61, CSIRO, Australia; Australian National University, Australia; Max Planck Institute for Biological Cybernetics, Germany

## Abstract

Adversarial examples are carefully crafted input patterns that are surprisingly poorly classified by artificial and/or natural neural networks. Here we examine adversarial vulnerabilities in the processes responsible for learning and choice in humans. Building upon recent recurrent neural network models of choice processes, we propose a general framework for generating adversarial opponents that can shape the choices of individuals in particular decision-making tasks towards the behavioural patterns desired by the adversary. We show the efficacy of the framework through two experiments involving action selection and response inhibition. We further investigate the strategy used by the adversary in order to gain insights into the vulnerabilities of human choice. The framework may find applications across behavioural sciences in helping detect and avoid flawed choice.

## Introduction

Advertisers, confidence tricksters, politicians and rogues of all varieties have long sought to manipulate our decision-making in their favour, against our own best interests. Doing this efficiently requires a characterization of the processes of human choice that makes good predictions across a wide range of potentially unusual inputs. It therefore constitutes an excellent test of our models of choice (Dan & Loewenstein, 2019). We have recently shown that recurrent neural network (RNN) models provide accurate, flexible and informative treatments of human decision-making (Dezfouli, Morris, Ramos, Dayan, & Balleine, 2018; Dezfouli, Ashtiani, et al., 2019; Dezfouli, Griffiths, Ramos, Dayan, & Balleine, 2019). However, how well these RNNs can interpolate and extrapolate outside the range of conventional inputs, and then the insights they can offer into human choice frailty, are unclear. To examine these issues, we require a systematic way of (a) finding the vulnerabilities in models of choice; and (b) proving that, and how, these vulnerabilities are also exhibited by humans. Here, we provide a general framework for doing this (Figure 1) based on RNNs fitted to behavioural data.

**Figure 1:**
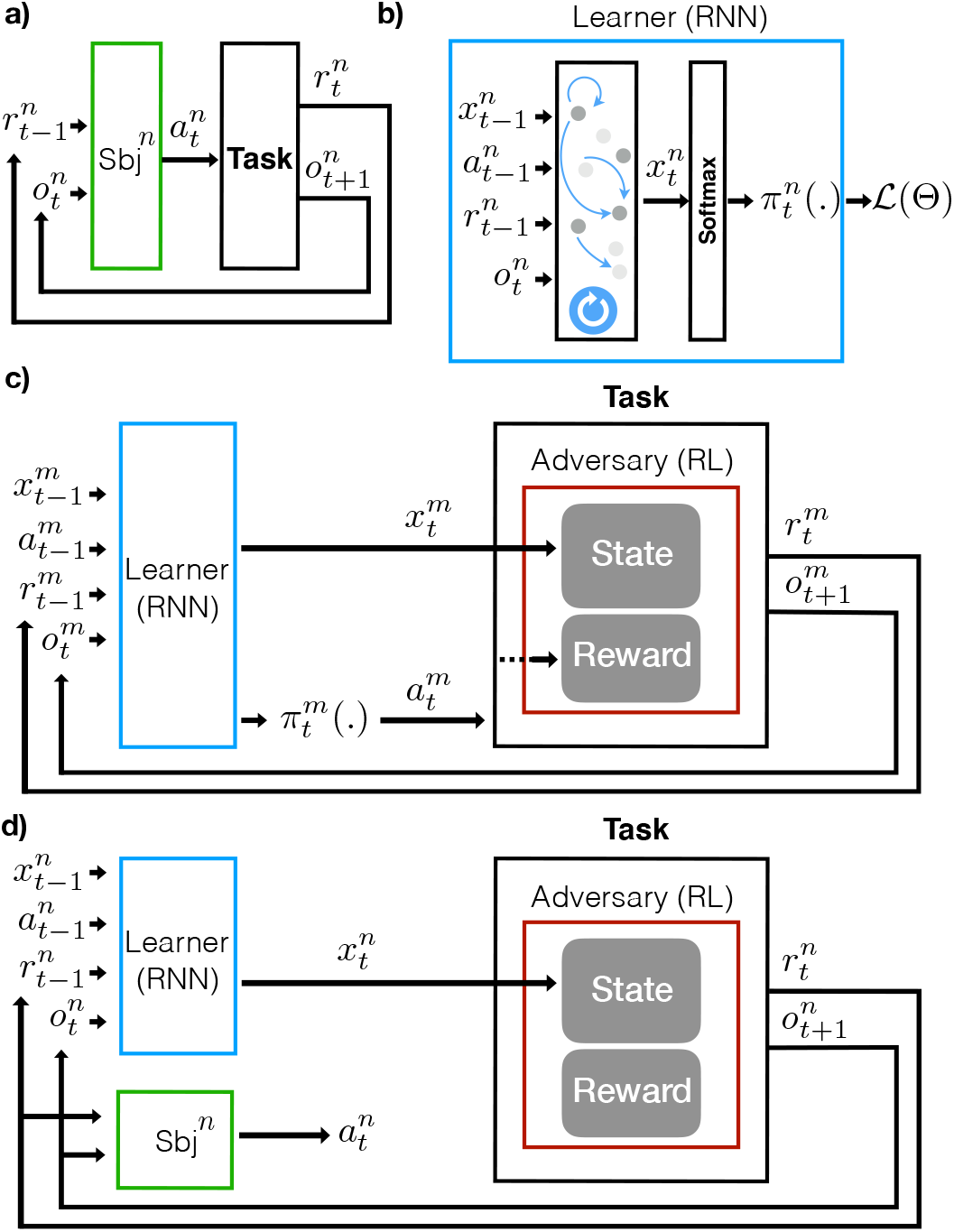
The framework. **a)** The interaction of the subjects with the task. At each trial *t*, subject *n* receives the learner reward 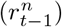 for its previous action and a new observation 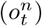 from the environment, and takes action 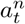. The environment then sends a new learner reward and observation back to the subject and this cycle continues until the end of the task. **b)** The behaviour of the subjects is modelled by a recurrent neural network (RNN; parameters Θ). The inputs to the RNN are the previous action 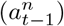, learner reward 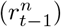 and the current observations from the task 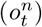 along with the previous internal state of the RNN 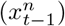. After receiving the inputs, the RNN updates its internal state, which is then mapped to a softmax layer to predict the next action 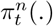. These predictions are then compared with the actual actions taken by the subjects to build loss function 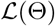, which is used for training the model. The trained model is called the learner model. **c)** The adversary is a reinforcement-learning (RL) agent which is trained to control the learner model. The internal state of the learner model (i.e., 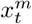 for simulation *m*) summarises its learning history, which is received by the dynamic adversary, determining its state (State). The agent then determines the learner reward and the next observation to be delivered to the learner model. The learner model subsequently takes its next actions and this cycle continues. If the action taken by the learner 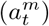 is aligned with the objective of the adversary, then the adversary receives an adversarial reward (Reward) which is used to train the adversary. Static adversaries are RL agents that do not have access to the internal state of the learner models and generates rewards and observations in an open-loop manner (not shown in the figure). Note that action 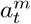 is *not* passed to the adversary, but is passed to the task to calculate the reward that should be delivered to the agent based on the reward that the adversary has assigned to each action. **d)** Using the trained adversary and the learner model for generating adversarial inputs to humans. Human subject n receives the learner reward for their actions and the observations from the trained adversary. They subsequently take an action 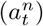 which is received by the learner model to update its internal state 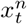. This state is then sent to the adversary to determine the learner reward for the action and the next observation. This cycle continues until the end of the task.

One line of systematic study of the first question started in image classification, with seminal early observations from Szegedy et al (Szegedy et al., 2013) that deep artificial neural networks are brittle to adversarial change in inputs that would otherwise be imperceptible to the human eye. This computer vision weakness of the machine has been an angle of attack to design adversaries for reinforcement-learning agents (Lin et al., 2017), followed by general formal insights on adversarial reinforcement learning on the more classical bandit settings (Jun, Li, Ma, & Zhu, 2018). To analyse human choice frailty, our framework involves two steps, the key one also involving a machine-vs-machine adversarial step in which a (deep) reinforcement-learning agent is trained to be an adversary to an RNN; this latter model is trained in a previous step to emulate human decisions following (Dezfouli et al., 2018; Dezfouli, Ashtiani, et al., 2019; Dezfouli, Griffiths, et al., 2019). This provides the general blueprint to tackling the first question. To show the promise of this framework to answer the second question, we applied it to two decision-making tasks involving choice engineering (Dan & Loewenstein, 2019) and response inhibition (Eisenberg et al., 2019) and tested the resulting adversaries on human subjects to assess the biases. We show that in both tasks, the framework was able automatically to specify adversarial inputs which were effective in steering choice processes to favour particular target actions. We further used simulations in order to illustrate and interpret the strategies used by the adversaries.

## The adversarial framework

The interaction of the subjects with the task is shown in Figure 1a. On each trial *t*, subject (*n*) receives what we call the learner reward 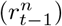 delivered by their previous action 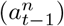 and observes the current state of the task (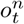; e.g., cues on a computer screen). They then take an action, 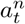. The process then repeats with the subjects receiving the learner reward of the action chosen 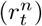 and the next observation (i.e., 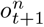).

In the non-adversarial case, learner rewards and observations are typically generated by a particular, pre-defined Markov decision process. For instance, actions might lead to learner rewards with certain fixed probabilities. By contrast, in our case, an adversarial model operates behind the scenes, deciding the learner reward of each action and the next observation that will be shown to the subject. To avoid triviality, adversaries work within budget constraints, which might limit or equalize per action the total number of learner rewards that the adversary could deliver. Within such constraints, the adversary faces a sequential decision-making problem to pick the learner rewards and observations that will shape subjects’ behaviours over the long run according to the target pattern, e.g., will lead them to prefer a certain action over the others or to take incorrect actions in certain situations.

We modelled the adversary using a reinforcement-learning (RL) agent. On each trial, the agent receives the learning history of the subject as input, and produces as output the learner reward and the next observation to be delivered to the subject. In RL terms, the learning history of the subjects constitutes the state of the agent, and its actions are the learner reward and the observation inputs for the subjects. The immediate adversarial reward for the agent reflects whether its output made the subjects meet the target behaviour. Within this structure, the adversary is trained to earn the maximum sum adversarial reward over the whole task, corresponding to producing outputs which most effectively push the subjects towards target actions.

In principle, the adversary could be trained through direct interactions with humans. In practice, however, this approach is infeasible given the delays of interacting with humans and the large number of training samples required. Instead, we use an alternative approach based on recent RNN models of human choice processes (Dezfouli et al., 2018; Dezfouli, Ashtiani, et al., 2019; Dezfouli, Griffiths, et al., 2019). Here, an RNN, called the *learner model* is trained from data collected from humans playing the task non-adversarially, and is then used to predict human behaviour under the different inputs that the adversary might try. RNNs are suitable because they provide a flexible differentiable family of models which are able to capture human choice processes in detail. Our approach has two additional advantages over training the adversary against humans: first, it can be carried out in settings where substantial training data (non-adversarial) already exists; second, our approach comes at the reduced human cost of just modelling the human behaviour for the non-adversarial task, which can then be used as input for diverse adversarial training scenarios.

### Learner model

In detail, the learner model (Figure 1b; determined by parameters Θ) comprises an RNN and a softmax layer which maps the RNN internal state to a probability of selecting each action (Dezfouli, Griffiths, et al., 2019). On trial *t* for subject *n*, the RNN layer has an internal state denoted by vector 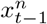, which reflects the RNN’s inputs up to that trial. This state is then updated on each trial based on the previous action 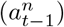 and learner reward 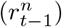, the current observations 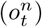, and, via recurrence, the internal state 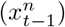. The output of the RNN layer 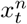 is passed to a softmax layer to predict the next action (i.e., π_t_(·)). The prediction is compared with the subject’s actual action 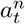, resulting in loss 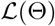 that is used for training the network. See Material and methods for more details.

### Training the adversary

The adversary is modelled as an agent which delivers learner rewards and observations on each trial, with the goal of maximising target behaviours subject to constraints. Since the agent’s choices on one trial can affect all the subsequent actions of the learner, the agent faces a sequential decision-making problem, which we address in an RL framework (Figure 1c).

The future actions of the learner after trial *t* depend on its prior history only through its internal state 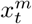 (for simulated learner *m*). Therefore, the RL agent uses 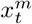 as the state of the environment to decide the learner reward 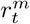 and next observation 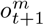 that it provides to the learner model. The process repeats with the new state of the learner model 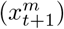 being passed to the adversary. Within this structure, the policy of the agent in choosing learner rewards and observations is trained to yield maximum cumulative adversarial reward subject to constraints. We used the advantage actor-critic method (A2C) (Mnih et al., 2016) and deep Q-learning (DQN) (Mnih et al., 2015) for training the RL agent. See Material and methods for more details on training and the constraints.

Along with this *dynamic* or closed-loop adversary, which is guided by the past actions of the subject (via the internal state of the RNN model), we also consider a *static* or open-loop adversary. This chooses learner rewards and observations without receiving the subject’s actions. Its policy is trained just as above, using samples generated from the learner model.

### Using the adversary

Figure 1d depicts how the trained dynamic adversary and the learner model are used in an experiment involving human subjects. The learner model does not choose actions, but receives the actions made by subject *n* as input, and tracks their learning history using 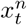. In turn, on trial *t*, 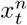 is received by the adversary to determine the learner reward 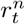 and the next observation 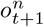 which the subject will use to choose their next action 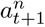. The same input, along with the actual action and learner reward, is delivered to the learner model. This cycle continues until the end of the task.

## Results

### Bandit task

This experiment is based on the bandit task introduced in (Dan & Loewenstein, 2019). On each trial, subjects make choices between two squares, one on the left of the screen, and the other on the right. After each choice, the subjects receive feedback about whether (smiley face) or not (sad face) their choice was given a learner reward. *A priori*, before each choice, the adversary assigns an outcome to both potential actions (e.g., that the left action will be rewarded and the right action will not get rewarded), and this is faithfully delivered *ex post* based on the subject’s choice. The goal of the adversary is to assign learner rewards to the actions in a way that makes subjects prefer one of the actions (called the ‘target’ action) – over the other one (the ‘non-target’ action). The target action is pre-determined (e.g., before the experiment starts the left action is set as the target action). The adversary is required to achieve this goal under a constraint: it must assign exactly 25 *a priori* learner rewards to each action, i.e., it cannot simply always assign learner rewards to the target action and no learner reward to the non-target action.

### Q-learning model

We first evaluated the framework in the synthetic setting of a *Q*-learning model. We generated data from a *Q*-learning model (1000 agents) with the same parameters used in (Dan & Loewenstein, 2019), and used these data to train the learner model. Then we trained the adversary RL agent to exploit the learner model. The agent received an adversarial reward every time the learner model chose the target action. The constraint was enforced at the task level, i.e., after the agent has allocated 25 learner rewards to an action, no more learner rewards will be allocated to that action. Conversely, if the agent has only assigned 25 — *k* learner rewards to an action by trial *T* — *k* (for *k* > 0, where *T* is the maximum trial number), that action will be assigned a learner reward on the remaining *k* trials.

The trained adversary was evaluated against both the learner model and the *Q*-learning model. The main dependent variable is the ‘bias’, which is the percentage of trials on which the target action was chosen (Figure 2a). As the figure shows, the adversary was able to guide the choices towards the target action. The average bias in playing the *Q*-learning model was 73.4% (ADV vs. QL column). This is similar to the results obtained in (Dan & Loewenstein, 2019), but here the results are obtained *without* knowing that the underlying model is *Q*-learning. The average bias when the adversary was simulated against the learner model is 73.8% (ADV vs. LRN column), which is comparable with the results against the *Q*-learning model.

**Figure 2:**
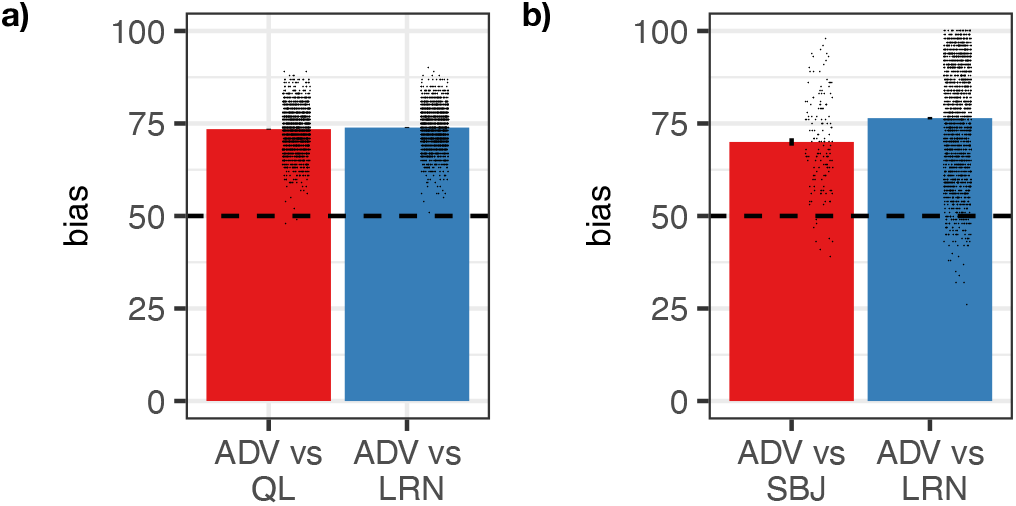
The performance of the adversary (’ADV’) in the bandit experiment against various opponents. ‘bias’ is the percentage of target actions selected by the opponent in response to learner rewards assigned by the adversary; error-bars represent 1SEM. Each black dot represents one simulation. **a)** ADV against an actual *Q*-learner (’QL’; red) or the learner model trained on *Q*-learners (‘LRN’; blue). **b)** ADV against human subjects (’SBJ’: red) or the learner model trained on other human subjects from (Dan & Loewenstein, 2019) (’LRN’: blue). Horizontal dashed lines represent equal selection of actions.

Next, we sought to uncover the strategy used by the adversary. Two 100 trial simulations are shown in Figure 3a (ADV vs. Q-learning). The blue and red circles indicate that the learner model selected the target and non-target action respectively. The vertical blue and red lines indicate that a learner reward was assigned to the target and non-target actions respectively. No line indicates that the learner reward was not assigned to the corresponding action. The green shaded area shows the probability the learner model awards to the target action. The general tactic used by the adversary is to assign a few learner rewards to the target action in the first half of the task; these few learner rewards, however, were sufficient to keep the probability of choosing the target action around %70-80 (shown by the green area), which is because the adversary never delivered non-target learner rewards in this period. Towards the end of the task, the adversary ‘burns’ the non-target learner rewards whenever the probability of the target action is above chance; at the same time the density of target learner rewards is increased to cancel the effect of non-target learner rewards. The combination of these strategies made the learner model choose the target action around 73% of the trials.

**Figure 3:**
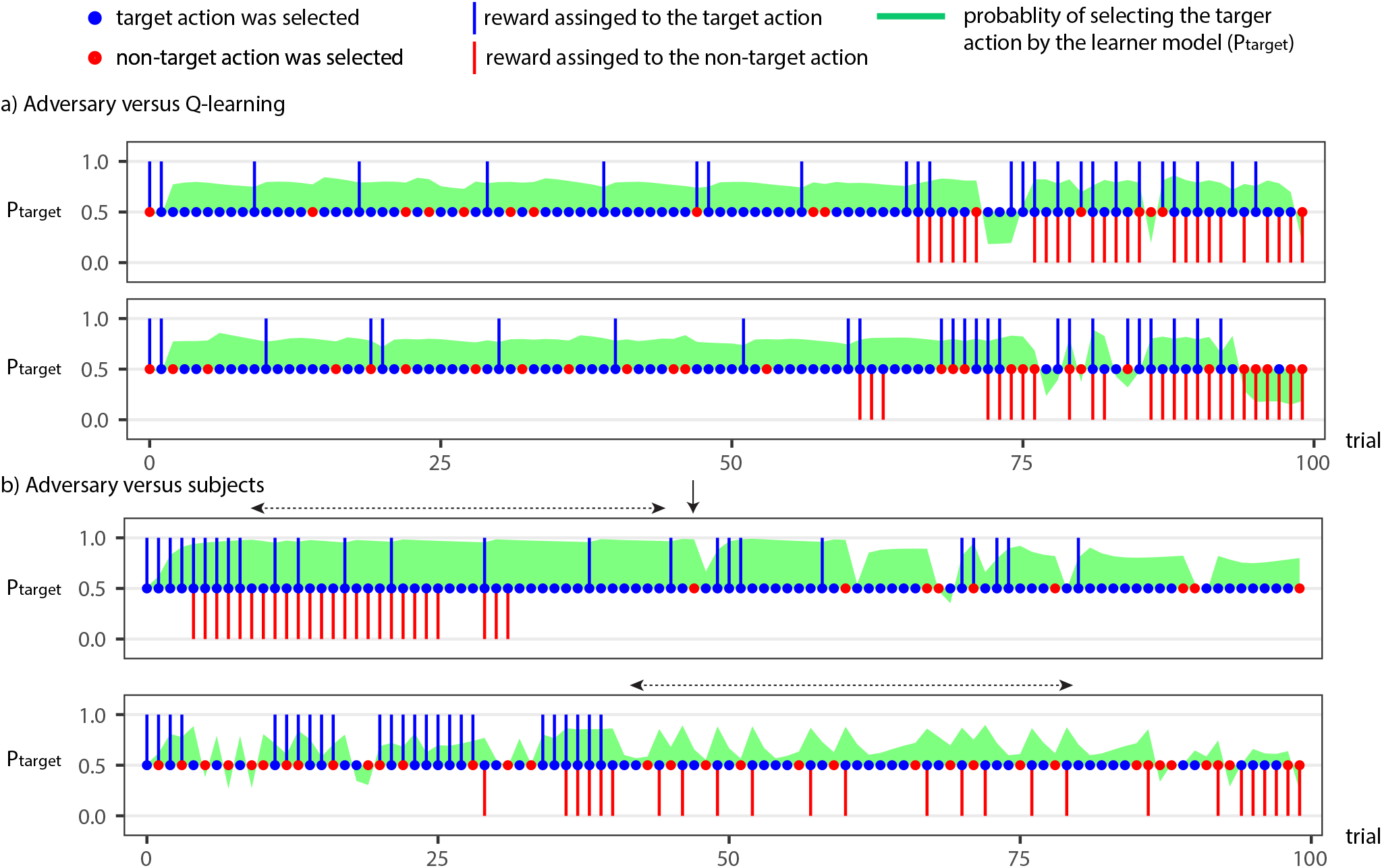
Simulations of the adversarial interactions over 100 trials in the bandit task. Blue and red circles indicate the selection of the target and non-target actions respectively. Vertical blue and red bars indicate the assignment of learner rewards to target and non-target actions respectively. The green area shows the probability the learner model accords to the target action. **a)** Two simulations of the adversary against a *Q*-learning model. **b)** Simulations of the adversary against the learner model trained using human behaviour. Solid and dashed arrows refer to the trials mentioned in the main text.

### Human subjects

We next applied the framework to develop an adversary for human choices in the task. The learner model was trained using the (non-adversarial) data published in (Dan & Loewenstein, 2019) for *N* = 484 subjects.

The trained adversary and the learner model were used to collect data from humans using Amazon Mechanical Turk (N = 157). The results are shown in Figure 2b. The bar ‘ADV vs SBJ’ shows the bias of subjects when playing against the adversary. The average bias was 70%, which is significantly higher than the equal selection of actions that might naively be implied by the equal numbers of learner rewards available for each choice (50% bias baseline; Wilcoxon signed-rank test, *p*–value ¡ 0.001). This shows that the adversary was successful in leading subjects to choose the target action. Figure 2b bar ‘ADV vs LRN’ shows the bias when the adversary is playing against the learner model (simulated). The average bias is 76.4%, which is better than when the adversary is pitted against human subjects. One major difference could be the subject populations used to train versus test the adversary.

To investigate the adversary’s strategy, we again simulated it against the human-trained learner model (Figure 3b). The adversary appears to seek to prevent subjects from experiencing the learner rewards assigned to the non-target action, but to make them see the learner rewards assigned to the target action, and therefore to select it. To achieve this, learner rewards are assigned to the target actions when this action is likely to get selected in the next trial, but for the non-target action, learner rewards are assigned when this action is *unlikely* to get selected.

Some of the tactics the adversary employs are evident in these simulations. In the top panel in Figure 3b, the adversary starts by continuously assigning learner rewards to the target action. Once the subject was set on choosing the target action, learner rewards are assigned to the non-target action to ‘burn’ them without subjects noticing. A second tactic is using partial reinforcement after the initial serial learner reward delivery on the target action (shown by the dashed horizontal line above the panel). This saves target learner rewards for later while not materially affecting choice probabilities. A third tactic is applied when the learner model takes a non-target action, as shown by the vertical arrow on the panel; here the adversary briefly increases target learner reward density to bring the subject back to choosing the target action.

The second simulation (bottom panel in Figure 3b) shows a more complex strategy which allows to ‘discretely’ spend the non-target rewards. In the period indicated by the horizontal dashed arrow, the learner tends to take a target action (blue circle) after each non-target action (the red circle). This pattern of behaviour is detected by the adversary and exploited by assigning a learner reward to each non-target action after each selection of the non-target action. This efficiently hides non-target learner rewards from the subjects. Altogether, such tactics substantially bias the subjects towards the target action.

Across the two experiments, it is evident that the strategy used against humans is quite different from the strategy used against *Q*-learning. Indeed, if we apply the human adversary against a *Q*-learning model, the average bias will be 55.2 and if we apply the adversary developed for *Q*-learning to humans (on a learner model trained using human data) the average bias will be 58.1 (Figure S1). These differ markedly from the biases reported when the adversaries play against their corresponding learner models.

### Go/No-go task

Our second experiment involved the go/no-go task as implemented in (Eisenberg et al., 2019). On each of 350 trials, subjects see either a go stimulus (e.g., a blue circle) or a no-go stimulus (e.g., an orange circle). Go stimuli are common (%90 of trials) and subjects are required to press the space bar in response to them. No-go stimuli are rare and require subjects to withhold responding. In the non-adversarial case, no-go stimuli are uniformly distributed across trials. By contrast, the adversary rearranges the no-go stimuli to encourage the largest number of mistakes (i.e., pressing space bar to no-go stimuli, or withholding responding to go stimuli), without changing the number of each stimulus type. Similar to the previous experiment, the constraint (exactly 90% of trials should be no-go) is enforced by fiat at the task level.

There are two differences between this experiment and the bandit experiments. First, the adversary determines observations (go vs. no-go) rather than learner rewards. Second, we considered an open-loop adversary – i.e., one that did not receive the state information 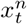 on trial *t*; and indeed knew nothing about what how the particular subject responded.

We started by training the learner model using the data generated in the random case (*N* = 770 subjects collected using Amazon Mechanical Turk were included in the analysis). Next, we trained the adversary using the learner model, but unlike the previous experiment, the adversary did not receive the state of the learner model. The trained adversary was then used to collect data from humans using Amazon Mechanical Turk (*N* = 139). The results are shown in Figure 4a. Subjects on average made 11.7 errors when playing against the adversary and 9.5 errors when no-go trials are distributed randomly (Wilcoxon rank-sum test; *p* < 0.001). Therefore the adversary was successful in finding a state distribution which induces at least a modest number of extra errors.

**Figure 4:**
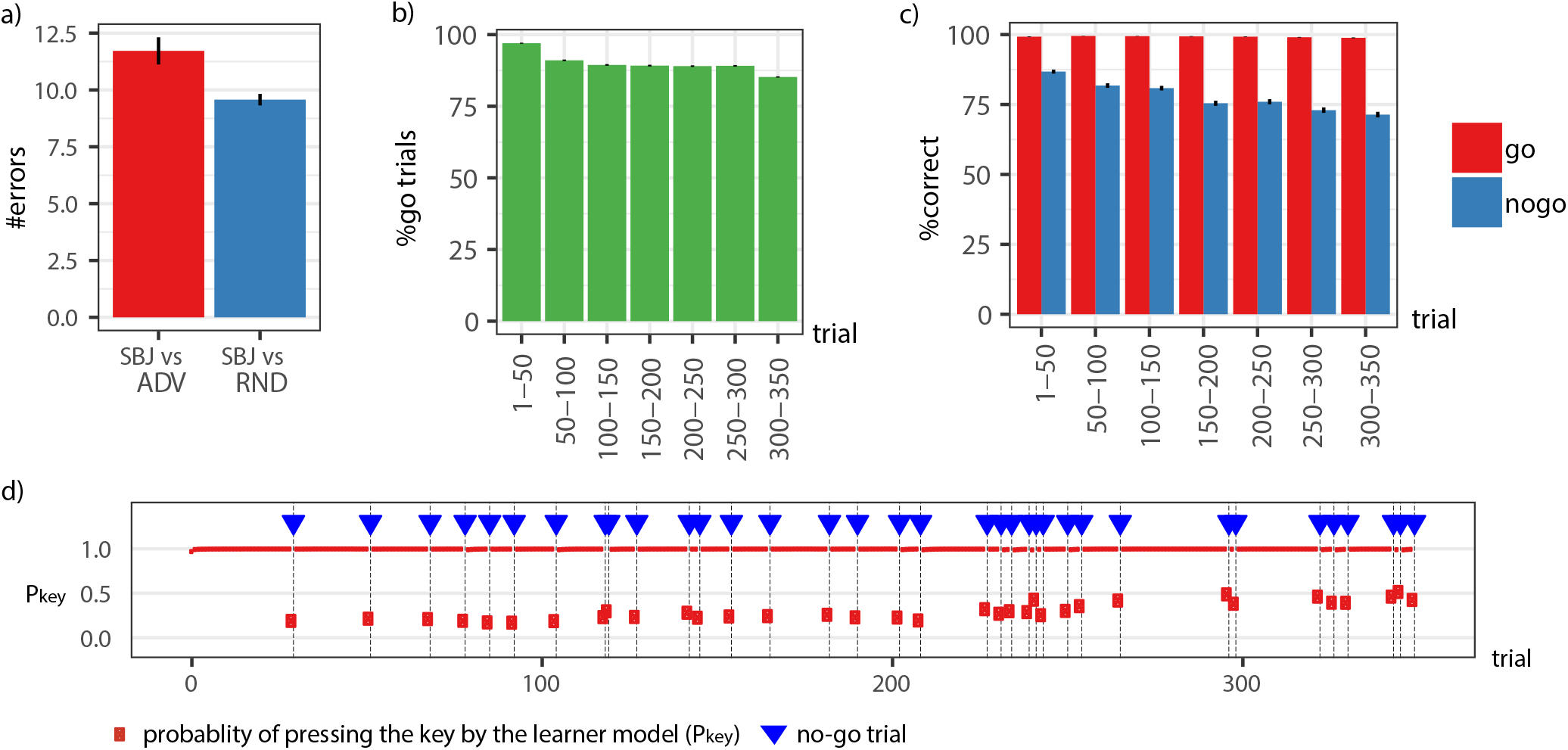
Adversarial strategy in the go/no-go task. **a)** The number of errors made by the subjects when no-go trials are delivered at random (‘SBJ vs ADV’) and when no-go trials are delivered by the adversary (‘SBJ vs ADV’). **b)** The distribution of the go trials over the task period delivered by the adversary. **c)** Percentage of the correct responses by the subjects in go and no-go trials when the no-go trials are delivered at random. **d)** A sample simulation. Blue triangles indicate no-go trials. The red shapes indicate the probability of pressing the key by the learner model. Error-bars represent 1SEM.

To elucidate the adversary’s strategy, Figure 4b shows the ratio of go trials across the task. The adversary allocates more no-go trials towards the end of the task. This may be since subjects (in the training data in which the no-go trials were randomly distributed) were more likely to make errors in the no-go condition later in the task (Figure 4c). However, it is a challenging task, since assigning all the no-go condition in a short period at the end of the task will likely have the opposite effect, since the subjects will have to stay alert only in a short period, and therefore a trade-off between these factors is required.

Figure 4d shows an example simulation. Blue triangles indicate no-go trials and red symbols indicate the probability of pressing the key (space bar) according to the learner model. Consistent with the data, the probability of making an error in the no-go trials increases over trials; thus the adversary spread the no-go trials across the task with a bias towards the end of the task to induce more errors.

## Discussion

We provided a general framework for generating adversarial models to elicit target behaviour in a wide variety of human choice processes. Two experiments showed that the framework is effective in inducing target behaviours and also interpretable using simulations.

### Cognitive biases

The strategies used by the adversarial models are driven by the choice characteristics embedded in the structure and weights of the learner model. Such choice characteristics can be seen as distilled, implicit, generalisations of the traditional cognitive biases (Tversky & Kahneman, 1974); and are what the adversary learns to exploit.

### Experiment design

Although we considered one-step adversarial scenarios, in principle the same framework can be used for multi-step experimental design (see also (Bak, Choi, Akrami, Witten, & Pillow, 2016)). Say, for example, that an experimenter desires the subjects to exhibit a specific pattern of behaviour, but the experimental parameters (e.g., probabilities, delays, etc) that yield the pattern are unknown. Following the framework here, the experimenter can train a learner model and use that model to train an RL agent to develop a static adversary which determines the optimal set of parameters for obtaining the desired behaviour. The obtained parameters then can be tested to see whether they make the subjects exhibit the desired behaviour; if they did not, the learner model can be re-trained using the new dataset and this process can be iterated until the desired behaviour is obtained. We conjecture that the procedure will often converge.

### Batch reinforcement-learning

An alternative to the framework developed here is using batch reinforcement-learning algorithms, which are able to learn from a pre-collected dataset without interacting with the environment (Lange, Gabel, & Riedmiller, 2012). One advantage of the current approach over these methods is that the policy of the adversary can be interpreted with respect to the behavioural of the learner model which has been used to train the adversary.

### Limitations

An assumption in the current framework is the generalisation of the predictions of the learner model from non-adversarial regimes to the adversarial regimes. The violation of this assumption implies that the adversary pushes the learner model in the parts of the state-space which have not been visited in the non-adversarial training and therefore the approximations of the human behaviour in those regions will be poor. This phenomenon is known as ‘extrapolation error’ in the batch reinforcement learning literature (Fujimoto, Van Hoof, & Meger, 2018) and several solutions have been suggested (see also (Cranko et al., 2019) for batch supervised learning), which can be applied to the current framework. Here, we used RNNs with a relatively small number of cells to avoid this issue, but the extension of the framework to address extrapolation error can be an interesting future step.

## Material and methods

### Training the learner model

The learner model was trained using the objective function: 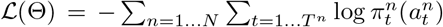, in which Θ refers to the free parameters of the model, *N* is the total number of subjects, and *T^n^* is the number of trials completed by subject *n*. Note that *π_t_*(*a_t_*) depends on the history of inputs, which are omitted for simplicity. Optimisation was based on the Adam optimiser (Kingma & Ba, 2014). The model was implemented using Tensorflow (Martin^~^Abadi et al., 2015) and the gradients were calculated using automatic differentiation. The recurrent neural network in the learner model was based on gated recurrent unit (GRU) architecture (Cho et al., 2014). The optimal number of training iterations (early-stopping) and the number of cells for each experiment were determined using 10-fold cross-validation. The optimal number of iterations and cells were then used to train a model using the whole dataset for each experiment. The number of cells considered was 5,8,10 cells, which was inspired by the previous works (Dezfouli, Griffiths, et al., 2019). The optimal number of cells and training iterations for each experiment is presented in Table S1.

### Training the adversary

The DQN algorithm was used for training the adversary in the bandit experiments (Mnih et al., 2015). The inputs to the RL agent were the internal state of the learner model (*x_t_*), its policy vector (probability of taking each action by the learner model), the trial number, and the of number rewards assigned to each action. Note that the internal state of the learner embeds other elements such as policy vector, but we also fed these elements explicitly to the RL agent to speed up the training. The output of the network was the value of each of the four actions corresponding to the combination of assigning reward/no-reward to each of the choices available to the learner model.

The model included three fully connected layers with 64, 64, and 4 units, with ReLU, ReLU, and linear activation functions. Models with replay buffer sizes of 200000, 400000 were considered. *ϵ*–greedy method was used for exploration with ϵ ∈ {0.01,0.1, 0.2}. Learning rates {10^−3^,10^−4^,10^−5^} were considered for training the model using Adam optimiser. These 18 combinations were trained and their performance was evaluated after {1, 2, 3, 4, 5, 6, 7, 8, 9} × 10^5^ training iterations. For the performance evaluation, the model was simulated against the learner model for 2000 times and the average bias was calculated. The model with the highest average bias was used. For the humans/*Q*-learning experiments, the highest bias was achieved with buffer size 400000/400000, ϵ = 0.1/0.01 and learning rate 0.0001/0.001

For the Go/No-go experiment, since the task was longer (350 trials), we used policy-gradient method and advantage actor-critic algorithm (A2C) (Mnih et al., 2016) for training the model. We also used an additional entropy term to encourage sufficient exploration (Ahmed, Le Roux, Norouzi, & Schuurmans, 2019). The input to the model was the current trial number and the total number of go/no-go states assigned. The output of the model was the policy (i.e., the probability of taking go/no-go). Note that the policy is stochastic. The network did not receive the learner internal state as input since we sought a static adversary. Both value and policy models had three layers, with 256, 256, and 1 unit(s) in the value layer, and 256, 256, and 2 units in the policy layer. The activation functions were ReLU, ReLU and linear respectively in each layer. We considered two values for the weight of entropy {0.01, 0.5} and the model with 0.01 entropy weight achieved a higher performance against the learner model (in terms of the average number of errors made by the learner model in 1500 simulation). The agents were implemented in Tensorflow and trained using Adam optimisation method (Kingma & Ba, 2014).

### Data collection

The study was approved by the CSIRO ethics committee (Ethics Clearance 102/19). The data were collected using Amazon Mechanical Turk. In both experiments, the participants received $0.4 USD. In the bandit task, they also could earn $0.01 USD for each smiley face.

For the bandit task, during the data collection, the adversarial model was running on a back-end Python server communicating with the task running in the web browser by receiving the action information from the task and sending the assigned rewards back to the test for the next trial. For the go/no-go experiment, since the delay between the trials was fast, it was not feasible to run the adversarial model on the back-end due to the communication lag. Instead, the adversarial models were exported to Javascript and ran in the web browser using Tensorflow.js framework.

For the case of the go/no-go task, we selected subjects with performance in the 75th percentile (which corresponded to less than 32 errors) and used their data for training the learner model. We then collected the data in the adversarial conditions and used the same threshold (less than 32 errors) to select the subjects who were included in the analysis. In the bandit task, all the subjects were included in the analysis.

## Acknowledgments

We are grateful to Yonatan Loewenstein for discussions. PD was funded by the Max Planck Society and the Humboldt Foundation.

## Supplementary Materials

### Task details

#### Bandit task

The implementation of the bandit task was based on the implementation in (Dan & Loewenstein, 2019):

We made the modifications to the task in order to communicate with a back-end for getting reward information before each choice. We also made modifications to the introduction pages of the task to be consistent with the ethics clearance requirements.

#### Go/no-go task

The implementation of the go/no-go task was based on the implementation in (Eisenberg et al., 2019):

We modified the code to interact with the adversarial model (using Tensorflow.js). We also made modifications to the introduction pages of the task to be consistent with the ethics clearance requirements.

---

**Figure S1:**
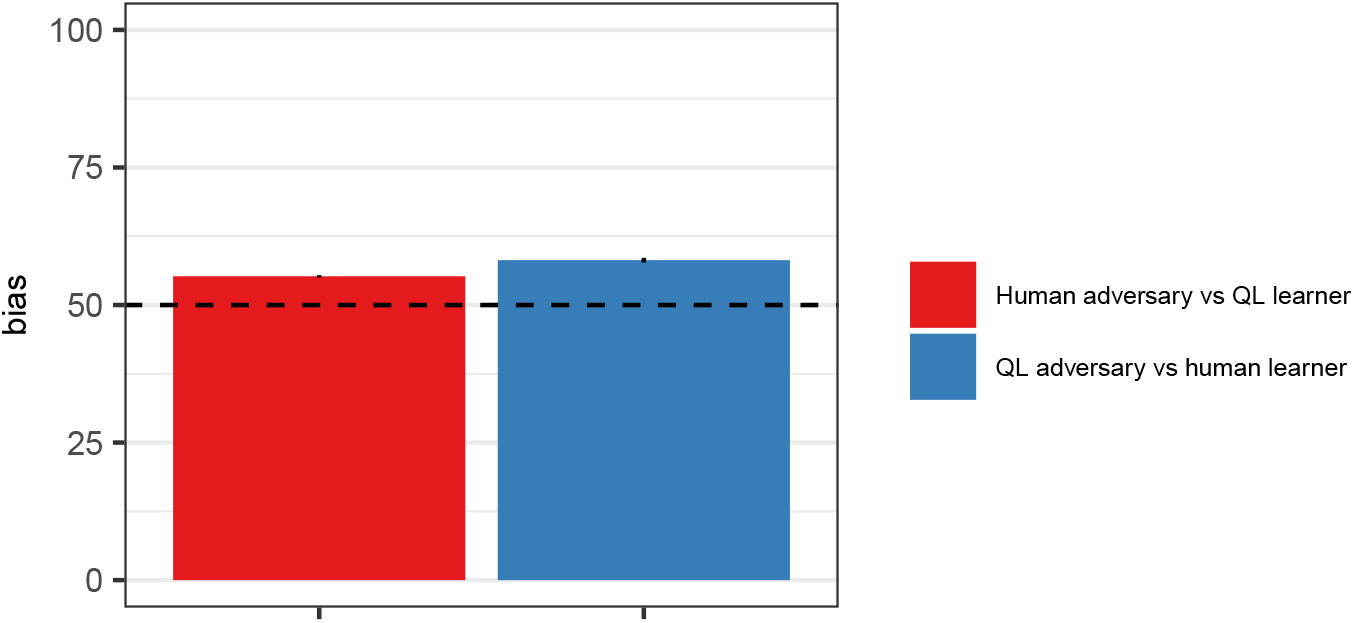
The performance of the human adversary when tested against the *Q*-learning learner model (Human adversary vs *QL*-learner), and the performance of the *Q*-learning adversary when tested against the human learner (*QL*-adversary vs human learner). Error-bars represent 1SEM.

**Table S1:** Optimal number of cells and training iterations.

| Experiment | #cells in RNN | #training iterations | learning rate |
| --- | --- | --- | --- |
| Bandit (*Q*-learning) | 5 | 46700 | 0.005 |
| Bandit (Humans) | 10 | 1100 | 0.005 |
| Go/No-go | 8 | 11600 | 0.001 |

